# Deep learning reveals the individual-level effects of artificial light at night on a wild insect

**DOI:** 10.64898/2026.06.15.732354

**Authors:** Ruonan Li, Rolando Rodríguez-Muñoz, Davide Dominoni, Tom Tregenza, Thomas O’Shea-Wheller

## Abstract

Artificial light at night (ALAN) is a widespread anthropogenic phenomenon with varied physiological, behavioural, and ecosystem-level effects. Its impacts have been studied extensively at the population level, however less is known about the individual changes that underpin these larger trends. We use a networked video system combined with GryllAI—a deep learning-based system for continuous individual monitoring—to explore this in the field cricket, *Gryllus campestris*. Applying field-realistic artificial light (10–25lx) or a control treatment to burrows, we continuously track the activity of 144 nymphs across >38,000h of video footage, recording life history outcomes for each individual. Results indicate that ALAN exposure does not influence daily activity timing, total activity duration, predation risk, or nymphal development duration. However, changes in fine-scale behaviour were apparent, with ALAN causing crickets to enter and exit burrows less frequently, especially at night; and spent greater proportion of time outside burrows during daytime. These behavioural adjustments were not evident from broad scale activity trends that could be observed manually. Consequently, our findings suggest that aggregate measures of activity may fail to capture the full scope of ALAN-mediated impacts in nature, and that automated monitoring techniques offer a promising means of addressing this.

## Introduction

Artificial light at night (ALAN) is an increasingly widespread consequence of urbanisation and other human activities [1, 2]. A growing body of research has demonstrated that ALAN can affect animal behaviour, physiology, and circadian regulation by modifying the perception of temporal cues in the environment [3, 4]. Because light is a primary cue for synchronising daily and seasonal rhythms, artificial illumination at night has the potential to disrupt the timing of biological processes even at relatively low intensities [5, 6]. Understanding how ALAN alters the temporal organisation of behaviour is therefore essential for assessing its ecological consequences.

A meta-analysis synthesizing studies across a wide range of taxa showed that exposure to ALAN can strongly affect physiological and behavioural processes, such as hormone levels, daily activity patterns, and life-history traits [7]. For instance, laboratory experiments on European perch (*Perca fluviatilis*) exposed to nocturnal light intensities of 0, 1, 10, and 100 lx found that melatonin production was suppressed even at 1 lx [8].

Similarly, a study assessing the effects of ALAN on the distribution and locomotor activity of a sandy beach isopod (*Tylos spinulosus*) in both the laboratory and field found reduced isopod abundance near light sources and a restricted tidal distribution range in the field, alongside decreased activity and disrupted circadian rhythms under ALAN in the laboratory [9]. In mammals, Swiss–Webster mice housed under either bright or dim light at night exhibit increased body mass and reduced glucose tolerance compared with mice maintained under a standard light: dark cycle, despite consuming a smaller proportion of food during the light phase, suggesting that even low levels of ALAN can disrupt the timing of food intake and metabolic signals, leading to excess weight gain [10]. Studies of birds also suggest that ALAN can alter circadian timing and extend perceived day length, leading to earlier activity onset, later activity offset, or increased overall activity [11–14]. ALAN induces earlier onset of dawn song and advances daily activity schedules in many songbird species, effects that have been suggested to confer adaptive reproductive advantages [15–18]. However, most direct evidence for such temporal shifts comes from birds, while existing work on insects has focused on when and how individuals move in response to artificial light, rather than on the timing and duration of activity. The timing of activity is a key determinant of ecological interactions and hence fitness, influencing exposure to predators, access to resources, and mating opportunities. In addition to substantial changes in total activity levels, animals may respond to altered light environments through fine-scale adjustments in movement and refuge-use behaviour, for example by modifying the frequency or duration of excursions from shelter.

Insects are sensitive to changes in light conditions, and numerous studies have shown that ALAN influences their behaviour, particularly through attraction to light sources and associated changes in movement and spatial distribution [1, 19, 20]. For instance, Van Langevelde, Ettema [21] demonstrated that lamps emitting shorter wavelengths attract higher abundances and species richness of moths. More recently, Charvalakis, van Boxem [22] showed that spectrum-dependent phototaxis in nocturnal insects varies not only among taxonomic groups but also over time, indicating that insect responses to artificial light have an important temporal component. Similarly, in an experiment involving 843 moths from 23 species, exposure to white LED lighting at 10 lx reduced activity by an average of 85% relative to natural night-time illumination. Amber lighting, often considered less disruptive, produced comparable effects at the same intensity [23]. In addition, artificial light pollution can interfere with celestial polarized skylight patterns [24], which are used for navigation by many insects [25–28].

Insect behaviour is often strongly structured in time, with activity typically concentrated within specific daily windows controlled by endogenous circadian clocks [29]. For example, studies of field crickets (*Gryllus campestris*) have demonstrated relatively stable individual chronotypes that are associated with fitness-related traits such as mating opportunities and predation risk [30]. In addition to individual variation, activity patterns may also differ between sexes, presumably because males and females experience different ecological and reproductive constraints that shape behavioural strategies [31–33]. Such differences can influence both the timing and duration of daily activity and therefore represent an important factor when examining behavioural responses to environmental change. Despite growing interest in the ecological effects of artificial light at night (ALAN), few studies have explicitly quantified how ALAN affects activity duration and activity timing in individual wild insects or examined the potential fitness-related behavioural consequences. As a result, the effects of ALAN on the timing and duration of insect activity, and associated fitness related behaviours under ecologically relevant conditions remain poorly understood.

The ‘WildCrickets’ system provides a useful framework for addressing these questions in nature, utilising networked cameras to record individual crickets (*Gryllus campestris*) [34–36]. To date however, the large quantities of data generated by this system have posed challenges for manual observation. Consequently, we have developed ‘GryllAI’, a fully-integrated deep learning pipeline that allows for the automated detection, tracking, and behavioural quantification of crickets filmed by the WildCrickets architecture. Here, we utilise this system to examine whether ALAN alters field cricket activity patterns and refuge use. Chronology in this species is closely linked to light conditions, which in turn may influence predation risk. We probe this question via automated extraction of individual activity durations and movement in and out of refugia. We explore whether changes in activity are associated with increased predation probability or altered refuge use during development. Our experiment utilises a relatively dim additional light source that is calibrated to mimic the light from local streetlamps.

We tested five predictions in pre-adult crickets immediately before the spring breeding season: 1) by extending the effective activity window, ALAN would alter activity timing, advancing onset, delaying offset, or both, thereby increasing activity during dusk, night and dawn but not daytime; 2) by altering perceived risk or environmental cues, ALAN would induce fine-scale behavioural adjustments, including changes in the frequency of burrow exits and returns; 3) ALAN would increase burrow abandonment, reflecting altered risk assessment or spatial behaviour; 4) ALAN would increase predation risk, because crickets may spend more time outside burrows or become more visible to predators; and 5) ALAN would accelerate emergence to adulthood, consistent with an extended photoperiod altering seasonal timing or increasing foraging opportunities.

## Methods

### Study system

Our study was carried out at the “WildCrickets” meadow near Gijón, northern Spain, which hosts a long-term monitored population of the field cricket *Gryllus campestris* (www.wildcrickets.org) [37]. This population has been continuously studied for more than two decades, providing extensive background information on individual life histories and behaviour. *G. campestris* is an annual species in which nymphs construct burrows early in development and overwinter in diapause before reappearing above ground in spring. [38, 39]. Nymphs and adults of both sexes are flightless and show strong site fidelity. Nymphs typically remain within a few centimetres of their burrow entrance. Burrows provide refuge from predators and adverse weather and form the spatial centre of most behavioural activity. The crickets in this study were the natural offspring of the previous generation. The only modifications to their natural environment were placing cameras beside studied burrows and lightly clipping grass near some entrances to keep them visible. Because a resident robin had learned to catch crickets at higher-than-usual rates in the previous breeding season, we placed wire mesh between each camera and burrow to reduce robin predation, although other predators such as shrews and lizards were not excluded. This did not eliminate robin predation, but it would likely have reduced it.

### Networked camera architecture

To facilitate continuous behavioural monitoring, we utilised a network of up to 140 cameras deployed across the ‘WildCrickets’ meadow, delivering 24h coverage of cricket activity. Each burrow was assigned a unique ID tag, and fitted with an infrared-capable camera (Vivotek IP8332) which recorded continuously at three frames per second. Cameras were connected to a centralised computer system running motion-detection software (iCatcher) so that frames were only captured when movement was detected at a burrow. Video data were recorded on 5 standard workstations using large capacity (8-16Tb) hard drives. At the end of the season these drives were physically transported to University of Exeter and uploaded to an external network-attached storage (NAS) server allowing for remote analysis.

### Automated monitoring pipeline

To automate the extraction of behavioural data from videos, we developed ‘GryllAI’, an end-to-end deep learning-based pipeline for remote cricket monitoring. This system is built around the YOLO11 (Jocher & Qiu, 2024) family of machine vision models, and utilises a hierarchical framework of detection and classification algorithms to extract and collate activity data directly from a NAS server. System development consisted of two key stages, building the central software architecture, and validating performance against manually collected data.

#### 1. System development

Fundamentally, the GryllAI system consists of a series of image detection models linked by a central data extraction and logic pipeline. This utilises separate models for cricket detection, burrow ID extraction, cricket orientation detection, cricket burrow status, and light level quantification, each being trained using images extracted from the NAS video archive. Across models, a total of 10,843 training images were used, sampled from the years 2013-2025, and selected to encompass the full range of biological and spatiotemporal variability present in the system. Training employed the PyTorch (Imambi et al., 2021) machine learning environment, with image augmentation to expand the quantity and diversity of data being provided via the ‘Albumentations’ (Buslaev et al., 2020) library. We opted to use the lightweight YOLO11n architecture specifically, as this provided a balance between processing speed and performance. Following training, models were integrated into a central data extraction and processing architecture developed in Python, allowing for detections to be recorded and synthesised into summary metrics via a graphical user interface.

#### 2. System operation

To analyse videos, the GryllAI pipeline extracts all individual frames from the NAS-hosted files and assigns date, time, and camera ID values derived from the file metadata. Frames are then passed sequentially to a central detection model, allowing for the identification and localisation of all crickets and burrow ID tags. Detected burrow ID tags are extracted, cropped, and rotated for analysis via an ID extraction model, yielding unique three-digit values linked to the corresponding frames. Extracted cricket images are then passed to an orientation model that detects the cricket’s head, rotates the image to standardise its orientation, and determines whether it is within or outside of the burrow. Across frames, a light level classification model simultaneously records whether the camera is in an IR or standard mode, thus inferring ambient light levels.

#### 3. Data extraction

GryllAI’s central processing pipeline cross-references and combines all model outputs into a single data row for each video frame. This details the camera ID, timestamp, cricket presence and count, cricket status within or outside of the burrow, and whether it is during the night or day. Data rows are then combined to produce a summary of daily activity for each burrow ID. Specifically, summaries partition total activity durations (defined as the cumulative time (in minutes) during which an individual cricket was observed outside of its burrow, even if only part of its body was outside of the burrow, and regardless of whether the individual was moving or stationary) into four time-windows: dawn (03:00–06:00), daytime (06:00–19:00), dusk (19:00–22:00), and night (22:00–03:00), and automatically calculate the beginning of activity time (BAT) and end of activity time (EAT) within each 24 hour period for any detected crickets. In addition, the duration and frequency of ‘Out’ events (defined as crickets being outside the burrow, even if only partially) were also summarised for each day across the four time-windows.

Extraction of these latter metrics is achieved via a simple set of processing rules, allowing GryllAI to search back through the history of detection events to determine when thresholds for each condition are met. Specifically, to calculate BAT, the system extracts the time of first cricket emergence from a burrow following the transition from night to day, ignoring any values that fall after 12:00. EAT is recorded as the last observed entrance into a burrow between 16:00 and 24:00, provided that the individual remains inside the burrow for at least the next 60 minutes. Individual ‘Out’ event durations are calculated as the time from when a cricket leaves the burrow (even partially) until it re-enters, thus recording each as a discrete period contributing to an overall count.

The definitions of BAT and EAT are conceptually consistent with those used in previous work [30], as both quantify daily activity onset and offset. However, in the present study, the night–day mode transition of the camera was used to identify the transition from night to day. This point is not affected by the artificial lights which are very dim compared to the sunrise. The earliest BAT recorded in the present experiment was 05:27 AM. In the previous study, BAT values occurring before 05:27 AM accounted for only 2.7% of the entire dataset. Consequently, the minor difference in BAT definition between studies is unlikely to materially affect inferences regarding relationships between BAT and other individual traits.

Beyond these rules, the system incorporates additional logic to ensure robust data synthesis while avoiding common errors. To confirm that BAT events represent sustained activity, crickets must emerge from the burrow for >2 minutes before the event is recorded, thus precluding cases where crickets briefly emerge for a single frame potentially to test external temperature conditions or respond to intruders. To accurately quantify ‘out’ events within each time window, any event overlapping two consecutive windows is counted as a single event and assigned to the window in which it began. Its duration, however, is partitioned between the two windows according to the time spent in each. Further, when combining data, burrow ID is filtered to include only the most frequently detected value across frames, avoiding errors caused by temporary tag occlusion. Finally, to alert users to hardware issues, the system produces warnings for any frame losses >60 minutes in length, these typically being associated with camera power outages, and automatically produces warnings for any overlapping metrics to maintain data quality.

#### 4. Performance validation

To validate system performance, we first assessed the accuracy of individual models via standard machine learning metrics. For detection models, these measures comprised precision (the proportion of detections that are correct), recall (the proportion of all true target crickets detected), and F1 score (a harmonic mean of precision and recall), while for classification models, top-1 accuracy (the proportion of classifications that are correct) was utilised. Each model was tested against unseen validation data and then retrained with additional images until it achieved an F1 score or top-1 accuracy ≥0.90, indicative of high detection and classification fidelity (Table S1).

Following this, we tested combined system performance when quantifying daily activity durations, BAT, and EAT across 32 test videos (each representing a single date), covering 10 unique burrows, with 2–7 days of data per burrow. Metrics were validated against manually-derived ground-truth data (obtained by the operator RL by manually reviewing videos and annotating activity timing and durations), ensuring that mean differences between manual and system-derived activity duration and BAT/EAT remained <10mins and <20mins respectively, with concordant intraclass correlation coefficients of 0.917 and 0.999 (Figure 1, Figure S1, Table S2).

**Figure 1.**
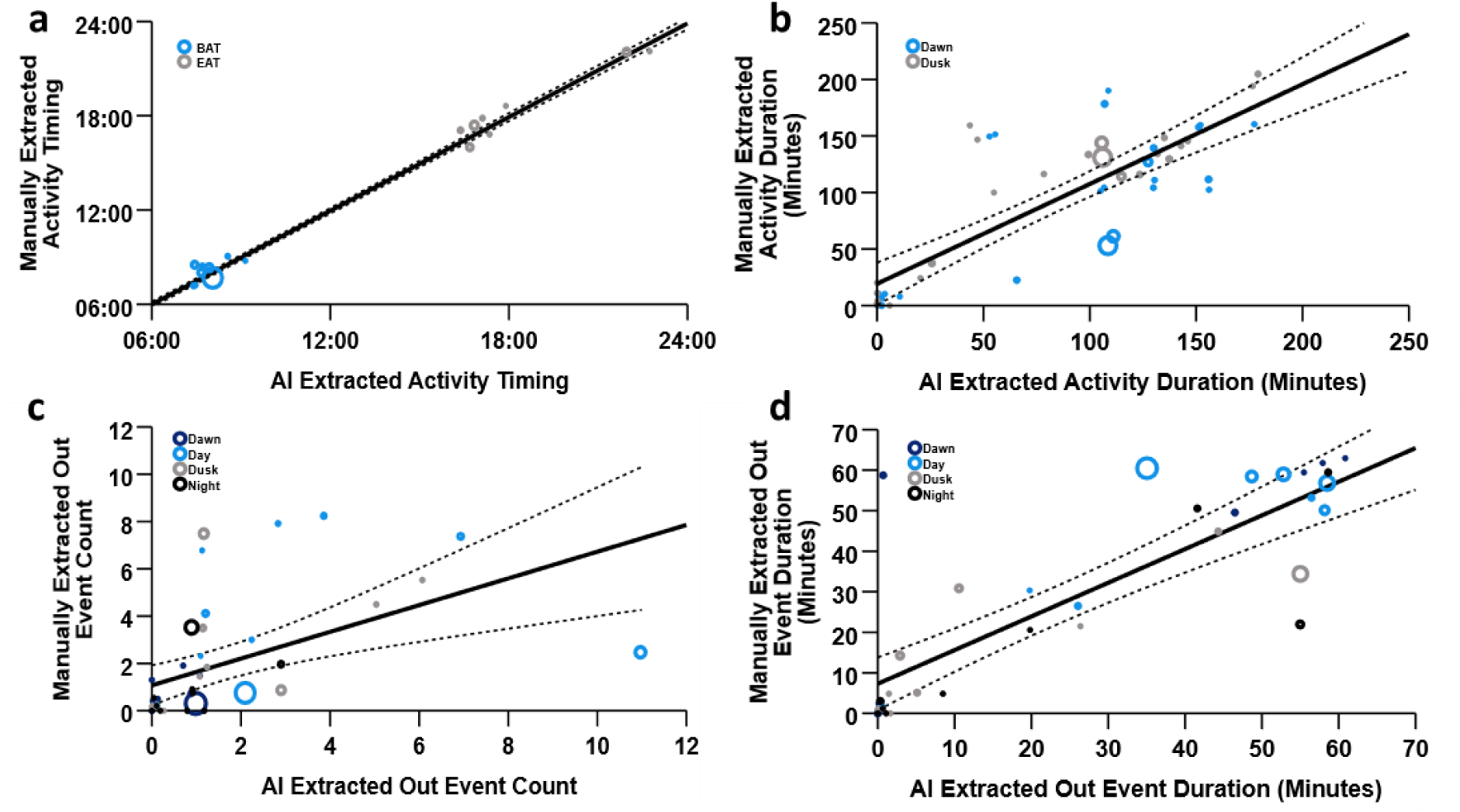
Comparison of automated and manually-derived results across system validation trials. (a, b) Correlation between AI and manually-recorded (a) BAT and EAT values; and (b) total dawn (03:00–06:00) and dusk (19:00–22:00) activity durations (N=32). Each point represents data from a single day and burrow combination, coloured by time window (dawn, blue; dusk, grey). Point size indicates the degree of difference between manual and AI derived values for (a) activity timing; and (b) activity duration within the same day and time window combination. (c, d) Correlation between AI and manually-derived (a) out event counts; and (b) out event durations (N=10). Each point denotes the value for a single burrow and day combination, coloured by time window (dawn, dark blue; day, light blue; dusk, grey; night, black). Point size indicates the degree of difference between manual and AI-derived values for (c) out event count; and (b) out event duration within the corresponding day and time window combination. Solid lines are linear trendlines fitted against the data, and dashed lines represent 95% confidence intervals.

We then validated individual ‘out’ event durations and counts using a further 10 videos (comprising one randomly selected date across the experiment period, used the same 10 unique burrows in the previous test), sampling a one hour period from each time window (00:00 – 01:00 for Night; 04:00 – 05:00 for Dawn; 12:00-13:00 for Day; 20:00-21:00 for Dusk). These were used to confirm that mean differences in activity event counts remained <2, while those of activity event durations were <11mins corresponding to intraclass correlation coefficients of 0.675 and 0.898 (Figure 1, Figure S2, Table S2).

### Experimental design

Field data were collected from late instar nymphs immediately prior to the 2025 breeding season. The experiment included 144 crickets: 80 individuals exposed to an artificial light at night (ALAN) treatment, and 64 individuals kept in natural night-time light conditions (control). At the start of the experiment, IR cameras were installed to monitor individual burrows and remained associated with a single cricket throughout the study. If a cricket abandoned its burrow, defined as leaving the burrow empty for more than 24 h, or was predated, the camera and, where applicable, light source were reassigned to a newly located nymph. The reassignment date was recorded as the start date for that individual.

ALAN was applied using a warm white surface-mounted LED (MULTICOMP PRO MCL053SWC YH1) with manually adjustable brightness (Figure S3). The LED was mounted on a metal stake beside the burrow, approximately 20 cm above the entrance, although this varied slightly with substrate conditions to maintain illumination within the target range (Figure S4). Illumination was adjusted to 10-25 lx using a handheld light meter. In total, 38,280 h of video footage were recorded and analysed using GryllAI, resulting in 2,134,322 data rows. To characterise activity, we extracted daily duration, event counts and duration of individual ‘out’ events for four time windows: dawn, daytime, dusk and night. BAT and EAT were automatically quantified for each individual over each 24h cycle. In addition to these metrics, we manually recorded sex, the occurrence of predation events, occasions when a cricket abandoned its burrow, and dates of maturity to adulthood.

### Data analysis

All analyses were conducted in R (version 4.4.2) and RStudio (version 2025.09.2 + 418). Linear mixed-effects models (LMMs) and generalized linear mixed-effects models (GLMMs) were fitted using the packages *lme4, lmerTest, and glmmTMB*. Survival analyses were performed using the *survival* package. Model diagnostics were assessed using *DHARMa*, by simulating residuals with the *simulateResiduals*() function and visually inspecting the residual distribution. The variance inflation factors (VIFs) were assessed using *vif*() function from the *car* package. Marginal and conditional R^2^ values for mixed models were calculated using *r.squaredGLMM*() function from the *MuMIn* package.

#### 1. Activity timing and activity duration

To test whether artificial light at night altered cricket activity, we analysed daily activity durations across the four time-windows of dawn (03:00–06:00), daytime (06:00–19:00), dusk (19:00–22:00), and night (22:00–03:00). Crickets remained inactive for 10.65% of all recorded observations across different windows, potentially reflecting behavioural decisions or other unmeasured factors.

To test whether ALAN affected activity duration when crickets were active, we fitted generalized linear mixed models (GLMMs) with a beta regression. The hourly proportion of activity duration was calculated from the hourly activity rate for each time-window record for each individual, representing the proportion of the 60-min interval during which the cricket was active. To meet the requirements of the beta distribution, zero values were replaced with a small constant (0.001), and 0.01 was added to the denominator in the proportion calculation to avoid proportion values of exactly 0 or 1. The hourly proportion of activity duration was used as the response variable, with light treatment, time window, their interaction, and sex included as fixed effects. Individual ID and the interaction between individual ID and date were included as random effects to account for repeated measures both within individuals and across days.

To examine whether light influenced activity in specific time-windows, we fitted separate models for each time window using the same model structure. The dataset of hourly proportions of activity duration was subset by the four time-windows, and each subset was analysed using the same model specification.

Daily activity timing was quantified using BAT and EAT values, converted into decimal hours. BAT was analysed using an LLM with light treatment and sex as fixed effects and individual ID as a random effect. EAT was analysed using a Gamma GLMM with a log link, again including light treatment and sex as fixed effects, and individual ID as a random effect. We also estimated the repeatability of BAT and EAT using the *rptR* package. BAT and log-transformed EAT were used as response variables in separate models, with sunrise or sunset included as fixed effects and individual ID fitted as a random effect.

#### 2. Fine-scale activity structure

There were 95,173 observations of ‘Out’ events in total. In the control group, each individual contributed an average of 787 observations across 12.6 days (SD = 518 observations; SD = 7.79 days). In the ALAN group, each individual contributed an average of 560 observations across 9.78 days (SD = 445 observations; SD = 7.08 days).

To assess whether ALAN altered the fine-scale structure of activity, we analysed both the frequency and duration of individual out events. Out frequency was modelled using a negative binomial mixed-effects model with log link, including treatment, time window, their interaction, and sex as fixed effects, with individual identity and date as random intercepts. Because time-windows differed in duration, log-transformed window length was included as an offset to account for unequal observation periods.

The duration of individual out duration was log-transformed and analysed using a linear mixed-effects model with the same fixed and random effects structure.

#### 3. Burrow abandonment

To test whether ALAN influenced the probability that an individual abandoned its burrow, we analysed the occurrence of these events using time-to-event analyses. For each individual, we recorded whether burrow-abandonment occurred during the monitoring period, and if so, when. A Cox proportional hazards model was fitted with time under monitoring as the time variable and burrow-abandonment as the event, including light treatment and sex as predictors. The assumptions of the Cox proportional hazards model were checked, and no violations were detected.

#### 4. Predation risk

To test whether exposure to ALAN increased predation risk, we again employed time-to-event analyses. For each individual, we recorded whether a predation event occurred during the monitoring period and the total number of days monitored. A Cox proportional hazards model was then fitted with predation as the event and monitoring duration as the time variable, with light treatment and sex included as explanatory variables. The assumptions of the Cox proportional hazards model were checked, and no violations were detected.

#### 5. Timing of maturity

To test whether ALAN affected the timing of emergence to adulthood, we conducted a time-to-event analysis using Cox proportional hazards models. As crickets were entered into the risk-set from different date, the start time for each individual was defined as the first day on which the individual was monitored, expressed relative to the baseline initiation date of the experiment (01/04/2025). End time was defined as the day of emergence, also expressed relative to the baseline day, or, for individuals that did not emerge, the final day of monitoring. Emergence status was treated as the event, and light treatment and sex were included as predictors.

## Results

### Effects of ALAN on timing of daily activity and activity duration

Both BAT (R = 0.11, CI [0.06, 0.16]) and EAT (R = 0.09, CI [0.04, 0.13]) showed significant repeatability within individuals, suggesting that crickets exhibit consistent daily activity timing. This pattern is consistent with our previous work [30], which also demonstrated that activity timing is repeatable within individuals across years.

We found no evidence that artificial light at night (ALAN) altered cricket activity timing. Neither the timing of activity onset (BAT) nor activity cessation (EAT) differed significantly between ALAN-exposed and control individuals (Table 1, Figure S5).

**Table 1.**
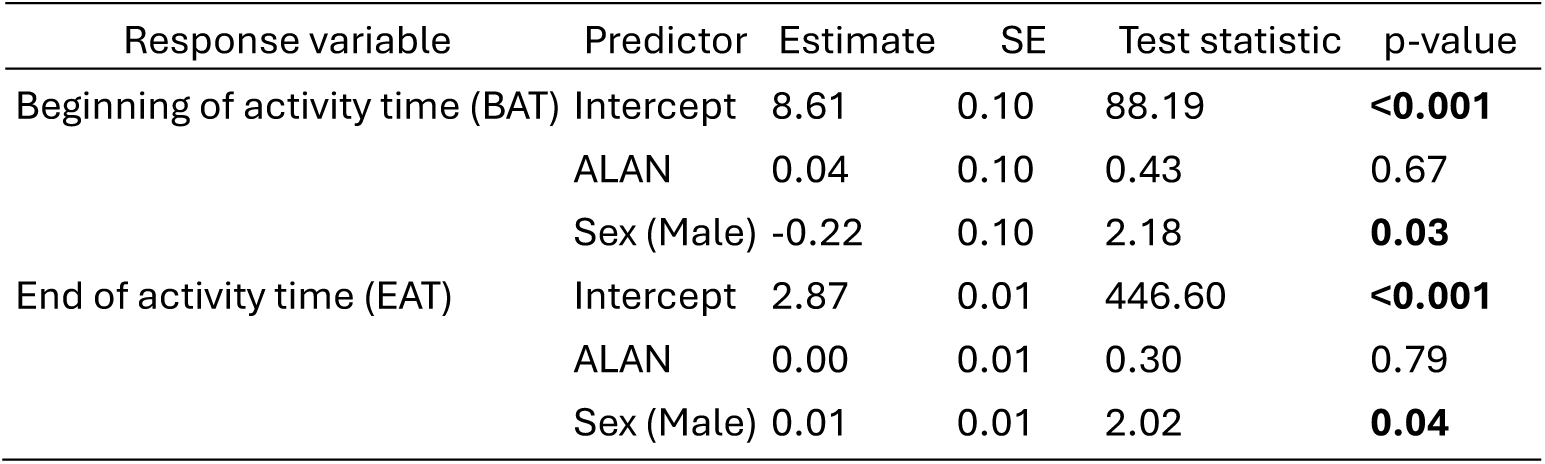
Effects of artificial light at night (ALAN) and sex on the beginning (BAT) and end (EAT) of activity time estimated from mixed-effects models.

Males and females differed in activity timing: males began activity earlier and ended it later compared to females (Table 1, Figure 2), indicating a longer daily activity window in males.

**Figure 2.**
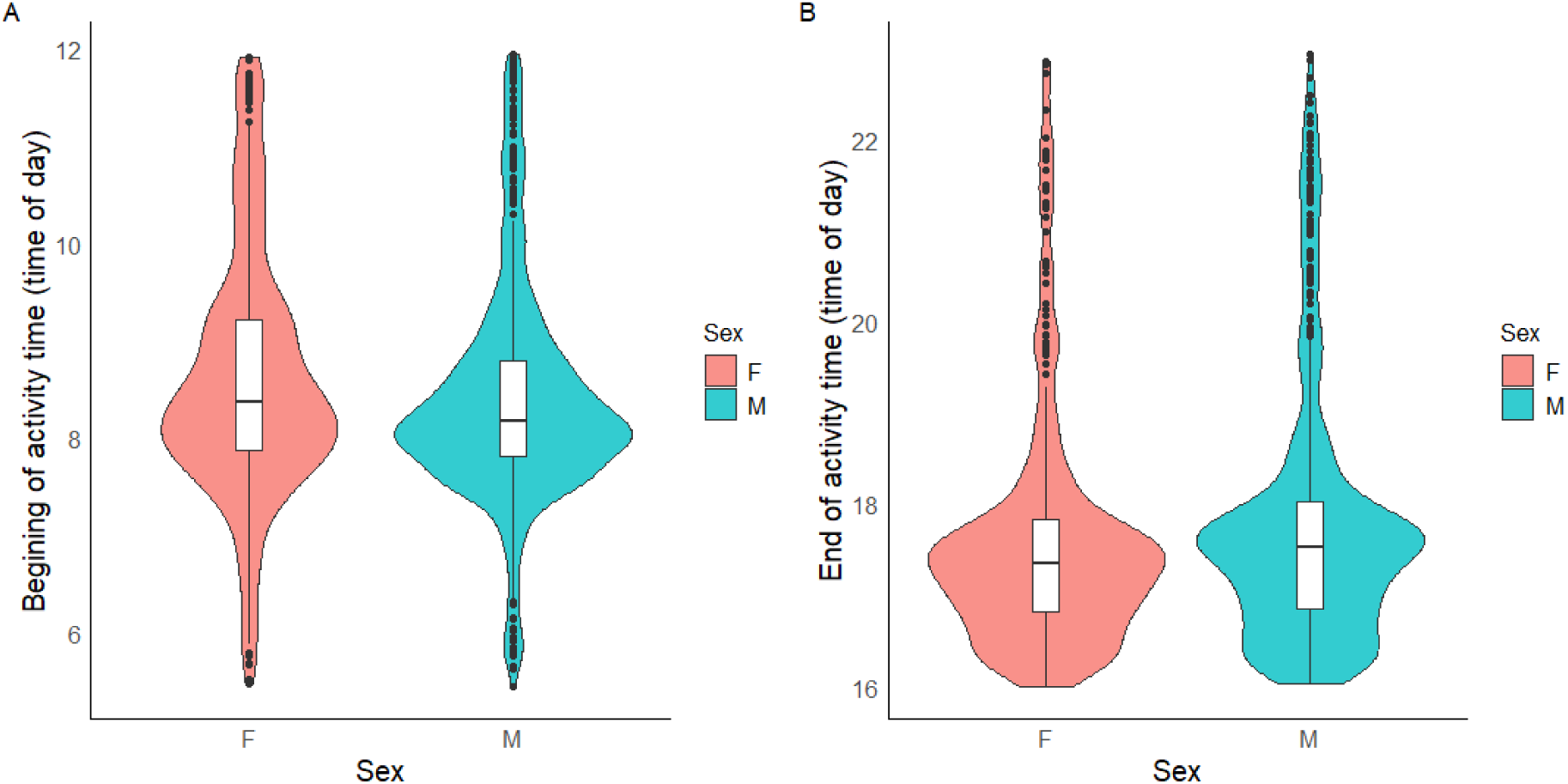
Sex differences in activity timings. Distribution of the beginning (A) and end (B) of activity time (hour) for female (F) and male (M) individuals. Violin plots represent the distribution density of the data, with embedded boxplots indicating the median and interquartile range. Points represent outliers beyond 1.5 × interquartile range.

ALAN exposure did not significantly affect cricket activity duration across the defined daily time-windows (Table 2). Activity duration did not differ between ALAN and control treatments in any of the four windows (dawn, day, dusk, or night) (Table 3). Males were active for a significantly greater proportion of the time than females during the daytime window (Table 3, Figure 3). No sex differences in activity were observed during the dawn, dusk, or night-time periods.

**Figure 3.**
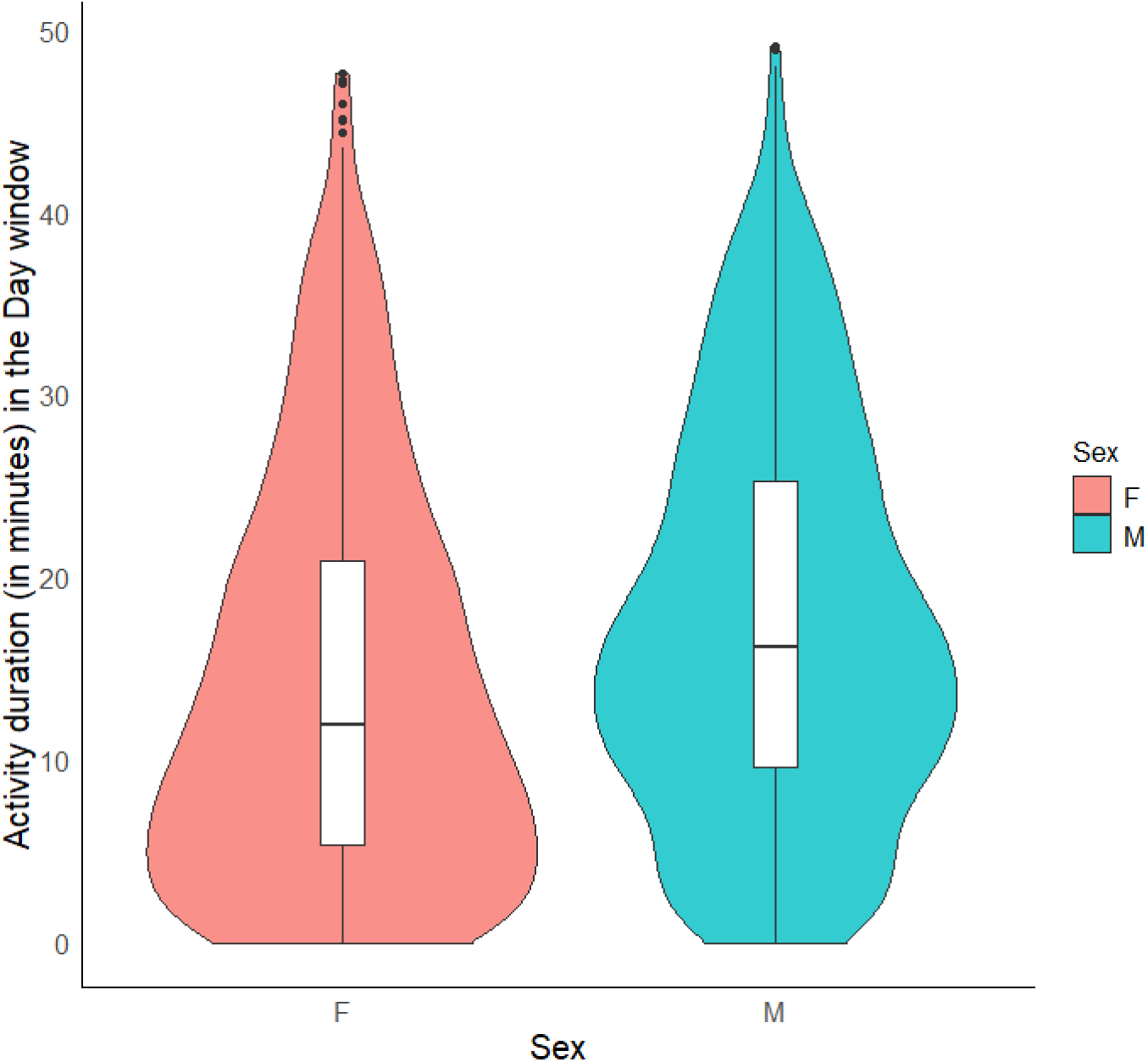
Sex differences of activity duration during the ‘Day’ window. Distribution of the activity duration (minute) for female (F) and male (M) individuals. Violin plots represent the distribution density of the data, with embedded boxplots indicating the median and interquartile range. Points represent outliers beyond 1.5 × interquartile range.

**Table 2.**
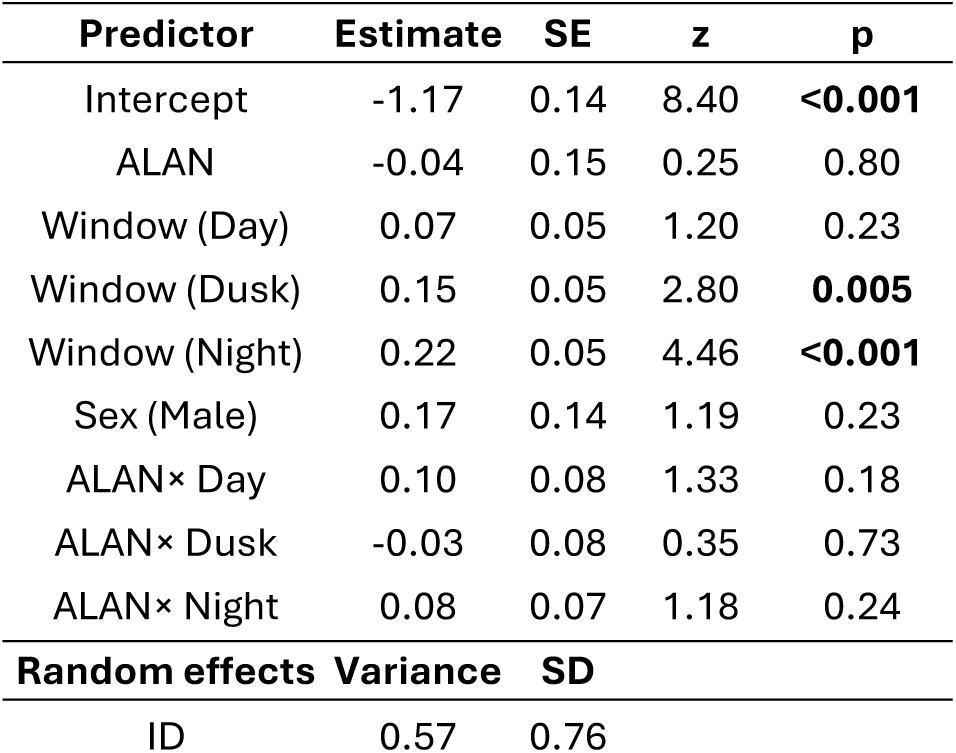
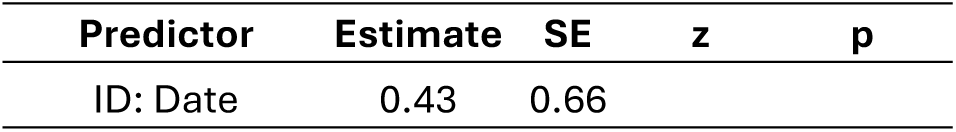
Effects of artificial light at night (ALAN), time window and sex on activity duration estimated from a general linear mixed-effects model with beta regression.

**Table 3.** Effects of artificial light at night (ALAN) and sex on activity duration across 4 different time-windows estimated from general linear mixed-effects models with beta regression.

| Window | Predictor | Estimate | SE | z | p |
| --- | --- | --- | --- | --- | --- |
| Dawn | Intercept | -0.97 | 0.14 | 6.91 | <0.001 |
|  | ALAN | 0.06 | 0.14 | 0.43 | 0.67 |
|  | Sex (Male) | 0.15 | 0.15 | 1.01 | 0.31 |
| Dusk | Intercept | -0.96 | 0.12 | 7.76 | <0.001 |
|  | ALAN | 0.02 | 0.13 | 0.13 | 0.90 |
|  | Sex (Male) | 0.16 | 0.13 | 1.26 | 0.21 |
| Day | Intercept | -1.37 | 0.09 | 15.35 | <0.001 |
|  | Sex (Male) | 0.40 | 0.12 | 3.35 | <0.001 |
| Night | Intercept | -0.85 | 0.16 | 5.35 | <0.001 |
|  | ALAN | 0.12 | 0.16 | 0.72 | 0.47 |
|  | Sex (Male) | 0.11 | 0.16 | 0.64 | 0.52 |

### Effects of ALAN on fine-scale Out event frequency and duration

To examine whether ALAN altered the fine-scale structure of activity, we analysed out event frequency (standardised by window duration) and the duration of time the cricket spent outside its burrow. Out frequency differed strongly among time-windows and was significantly modified by ALAN in a window-dependent manner (ALAN × Window interactions, all p < 0.001; Table 4, Figure 4). Overall, ALAN individuals made fewer out events, with the strongest reduction occurring at night.

**Figure 4.**
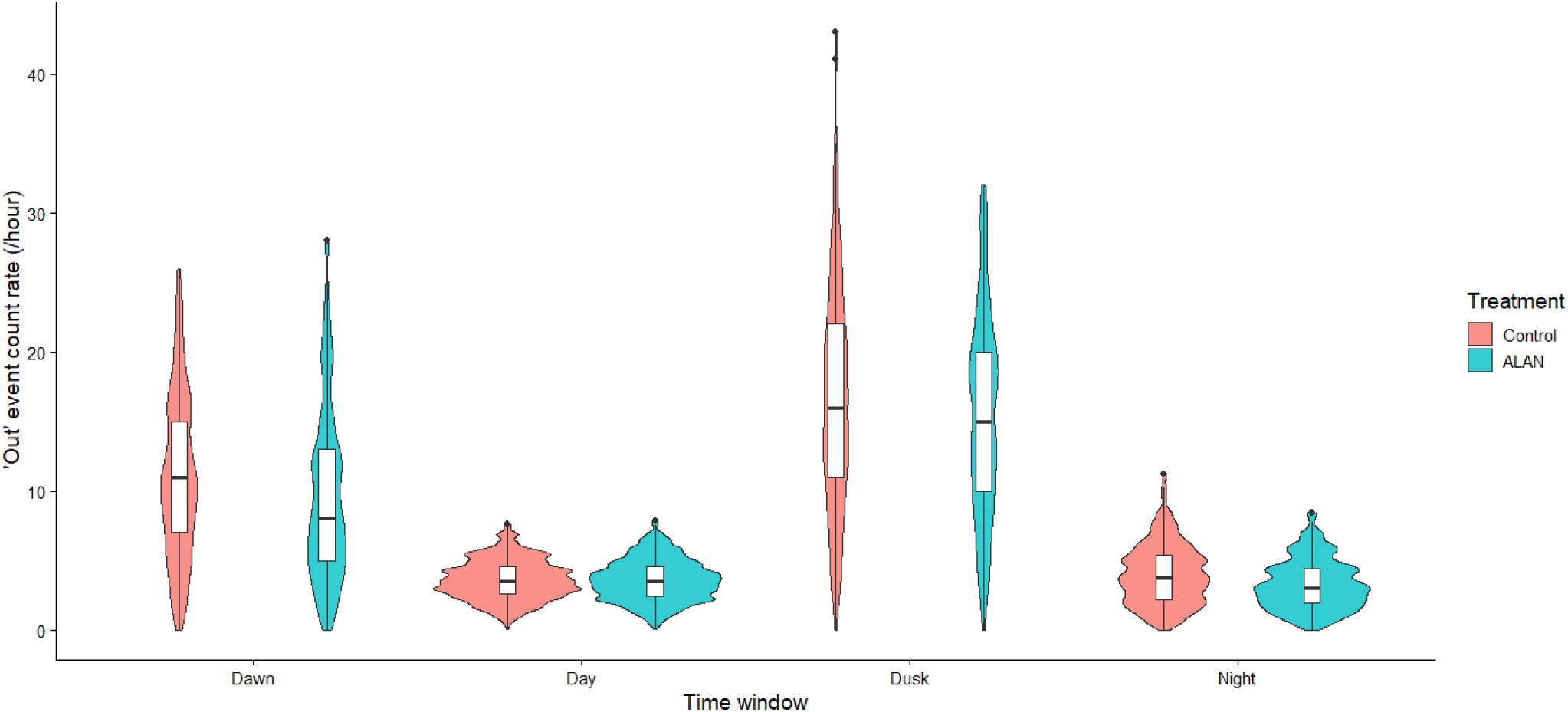
Distribution of ‘Out’ event frequency across time-windows under control and ALAN treatments. Violin plots show the distribution of outing rate (events h⁻¹) within each time window (Dawn, Day, Dusk, Night), with embedded boxplots indicating the median and interquartile range. Outing frequency varied across windows and was generally reduced under ALAN, particularly during the night.

**Table 4.** Negative binomial mixed model for Out event counting rate (offset by corresponding window duration)

| Term | Estimate | SE | z value | p value |
| --- | --- | --- | --- | --- |
| Intercept | 1.07 | 0.07 | 16.02 | **<0.001** |
| ALAN | -0.12 | 0.05 | 2.68 | **0.007** |
| Window (Day) | 0.02 | 0.01 | 2.64 | **0.008** |
| Window (Dusk) | 0.41 | 0.01 | 46.75 | **<0.001** |
| Window (Night) | 0.08 | 0.01 | 9.80 | **<0.001** |
| Sex (Male) | 0.01 | 0.04 | 0.18 | 0.856 |
| ALAN × Window (Day) | 0.07 | 0.01 | 6.48 | **<0.001** |
| ALAN × Window (Dusk) | 0.05 | 0.01 | 3.82 | **<0.001** |
| ALAN × Window (Night) | -0.07 | 0.01 | 5.19 | **<0.001** |

By contrast, ALAN had no overall effect on the time a cricket spent outside its burrow (out duration) (p = 0.585). Out duration varied across windows, and ALAN individuals showed slightly longer daytime out duration than controls (ALAN × Window (Day): p = 0.0068; Table 5, Figure 5), with no treatment differences during dusk or night.

**Figure 5.**
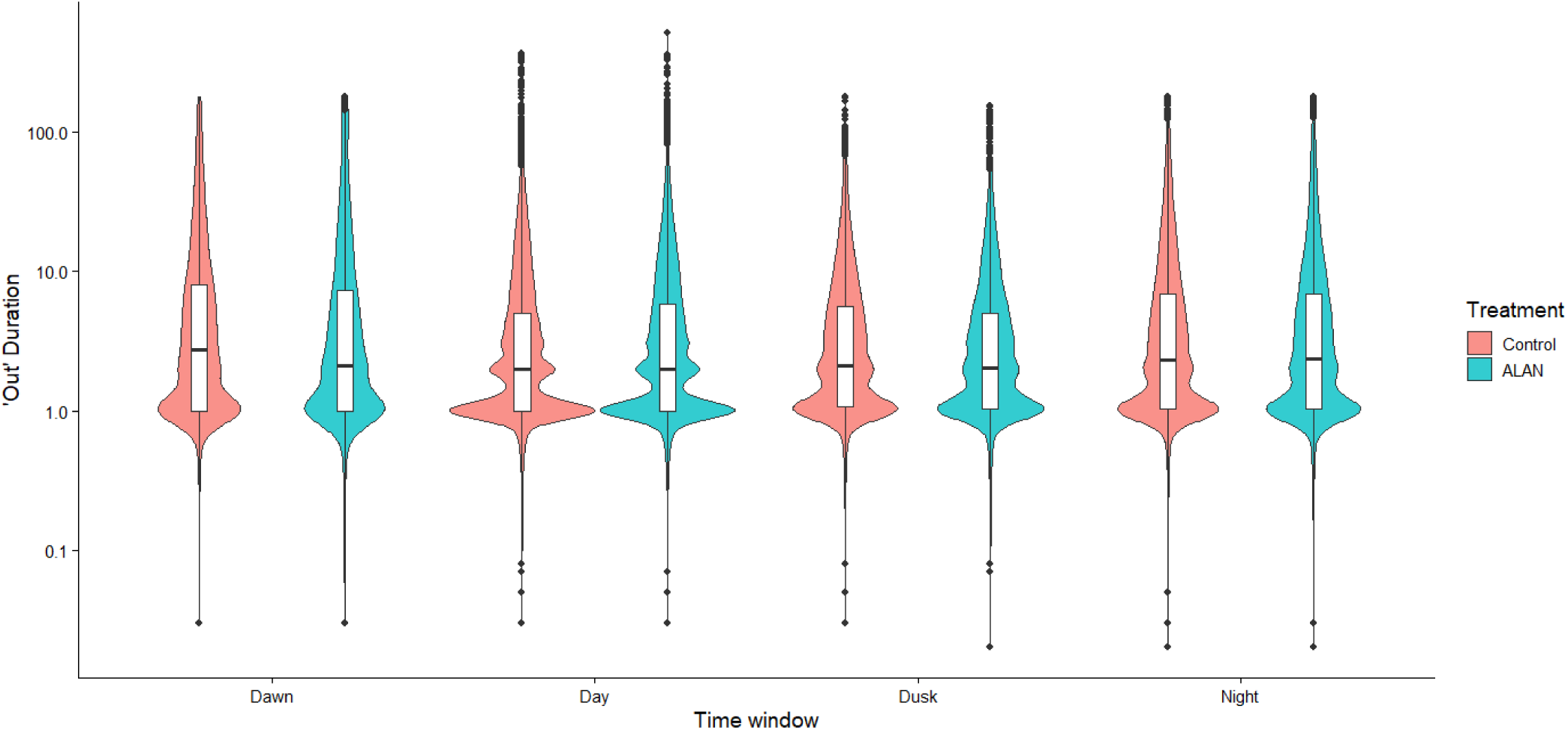
Distribution of ‘Out’ duration across time-windows under control and ALAN treatments. Violin plots show the distribution of individual out durations during each time window (Dawn, Day, Dusk, Night), with embedded boxplots indicating the median and interquartile range. Durations are presented on a log-scaled y-axis. ALAN individuals exhibited longer daytime outings than controls.

**Table 5.** Linear mixed model for log-transferred Out duration.

| Term | Estimate | SE | df | t value | p value |
| --- | --- | --- | --- | --- | --- |
| Intercept | 1.31 | 0.05 | 1522 | 24.43 | **<0.001** |
| ALAN | -0.03 | 0.05 | 24050 | 0.55 | 0.585 |
| Window (Day) | -0.33 | 0.02 | 95120 | 17.44 | **<0.001** |
| Window (Dusk) | -0.23 | 0.02 | 95050 | 10.24 | **<0.001** |
| Window (Night) | -0.09 | 0.02 | 95030 | 4.32 | **<0.001** |
| Sex (Male) | 0.04 | 0.04 | 1362 | 0.92 | 0.359 |
| ALAN × Window (Day) | 0.08 | 0.03 | 95130 | 2.71 | **0.0068** |
| ALAN × Window (Dusk) | -0.01 | 0.03 | 95060 | 0.36 | 0.719 |
| ALAN × Window (Night) | 0.05 | 0.03 | 95040 | 1.54 | 0.123 |

### Effects of ALAN on burrow-abandonment and predation risk

ALAN had no significant effect on predation risk, with no difference in the probability of predation between ALAN-treated and control individuals (Cox proportional hazards model: hazard ratio = 1.52, 95% CI = 0.62–3.72, p = 0.36; Table 6). Burrow abandonment was slightly more likely under ALAN, although this effect was weak and not statistically significant (hazard ratio = 1.74, 95% CI = 0.92–3.30, p = 0.09; Table 6).

**Table 6.** Effects of artificial light at night (ALAN) and sex on predation risk and burrow abandonment estimated from Cox proportional hazards models.

| Response | Predictor | Hazard ratio | SE | z | p | 95% CI |
| --- | --- | --- | --- | --- | --- | --- |
| Predation risk | ALAN | 1.52 | 0.46 | 0.92 | 0.36 | 0.62–3.72 |
|  | Sex (Male) | 1.17 | 0.45 | 0.35 | 0.72 | 0.49–2.82 |
| Burrow abandonment | ALAN | 1.74 | 0.33 | 1.69 | 0.09 | 0.92–3.30 |
|  | Sex (Male) | 1.23 | 0.32 | 0.66 | 0.51 | 0.66–2.29 |

### Effects of ALAN on emergence timing

ALAN did not significantly affect emergence timing, with days to adult emergence being similar between ALAN and control individuals (Table 7, Figure S6).

**Table 7.** Effects of artificial light at night (ALAN) and sex on days to emergence estimated from a Cox proportional h<u>azards model.</u>.

| Predictor | Hazard ratio | SE | z | p | 95% CI |
| --- | --- | --- | --- | --- | --- |
| ALAN | 0.99 | 0.26 | -0.04 | 0.97 | 0.59–1.66 |
| Sex (Male) | 1.36 | 0.27 | 1.12 | 0.26 | 0.80–2.31 |

## Discussion

We found no evidence that artificial light at night (ALAN) altered daily activity timings or durations in the field cricket *Gryllus campestris*. Individuals exposed to ALAN did not differ from controls in activity onset, activity cessation, or total activity duration across the four measured time-windows. Similarly, ALAN exposure had no detectable effect on life-history outcomes that might be affected by increased light: predation risk did not differ significantly in the increased light group, and ALAN did not accelerate maturity to adulthood. The mesh we used to reduce predation by robins (which are exclusively diurnal predators) may have reduced the effect that changes in daytime activity had on predation risk (Figure S3). However, many meadows occupied by crickets do not have resident robins, so this modification would simply make our experiment comparable to a site with lower levels of robin activity. ALAN-exposed individuals came out of their burrows less frequently, particularly at night, and spent longer periods outside during the day. This suggests that rather than producing overall shifts in daily schedules, ALAN appeared to affect the microstructure of behaviour in wild crickets.

One explanation for the limited effects of ALAN on broad activity schedules is that the burrow ecology of field crickets may buffer stronger behavioural disruption under ecologically realistic conditions. Unlike many flying nocturnal insects that are strongly attracted to light sources, *G. campestris* is a ground-dwelling species that spends a substantial proportion of time within a burrow that serves as a refuge from predators and environmental stressors. Artificial light applied at the burrow entrance may therefore result in relatively low effective illumination within the refuge itself. As a consequence, individuals may remain partly shielded from artificial illumination during periods of rest. Under these conditions, crickets may still detect ALAN, but the refuge environment may reduce its influence on broader activity timing, allowing daily schedules to remain similar to those observed under natural light conditions.

The ecology and life history of field crickets may further buffer them against broad behavioural disruption by artificial light. In contrast to highly mobile insects that rely on visual navigation or phototaxis, cricket nymphs typically remain within a radius of a 10-20 cm centred on their burrow. Activity primarily consists of short excursions around the burrow entrance rather than extensive movements across the landscape. Because their behaviour is tightly linked to their burrow, the temporal organisation of activity may be governed more strongly by internal rhythms and local microhabitat conditions than by ambient illumination. This behavioural ecology may therefore reduce the potential for artificial night lighting to substantially alter daily activity schedules.

Another explanation is that the intensity and duration of ALAN exposure in our experiment may have been insufficient to generate broad behavioural changes. Although we measured illumination at ground level from nearby streetlights at the field site and maintained comparable illumination levels in the ALAN treatment, several experimental studies showing strong biological effects of artificial light have involved relatively high illumination levels or prolonged exposure periods. For instance, a study on the leaf beetle (*Phaedon cochleariae*) found that time until adult eclosion was significantly prolonged only under continuous exposure to relatively high light intensity (50lx compared with 10lx) [40]. Similarly, in fruit flies (*Drosophila jambulina*), prolonged development and arrhythmic eclosion occurred only under high-intensity ALAN (50 lx) [41]. Crickets in our experiment experienced artificial illumination for an average of approximately 16 days during development. This relatively short exposure period may not have been sufficient for coarse-scale behavioural effects to emerge. Large-scale field experiments investigating artificial light at night have emphasised the importance of long-term exposure when assessing ecological consequences of illumination. For example, a five-year long-term experiment on macro-moth populations found that moth populations were negatively affected by the presence of light at night, but such effects only became apparent after multiple years of exposure [42].

Nevertheless, the reduction in the frequency at which crickets moved in and out of their burrows indicates that wild crickets were able to detect and respond to artificial illumination, even at relatively low light levels. Importantly, this response did not manifest as a shift in overall daily schedules, but as a change in behavioural microstructure. Individuals under ALAN appeared to leave and enter their burrow less frequently, particularly at night, while the time they spent outside their burrow during the day increased. This pattern may reflect more cautious excursions, increased reliance on the refuge, or a redistribution of activity across fewer but longer outings.

Behavioural adjustment is often the first response of organisms to environmental change, and can allow individuals to cope with altered conditions through a range of behavioural strategies [43, 44]. In some cases, environmental change leads to substantial shifts in overall activity schedules [45, 46]. However, in other systems, animals may instead respond through more subtle behavioural adjustments without large changes in overall activity timing. For example, lactating grey seals (*Halichoerus grypus*) can adjust fine-scale activity budgets by shifting time between resting and vigilance behaviours without altering overall daily activity patterns [47]. Our findings are consistent with this latter scenario, where ALAN modified the fine-scale structure of behaviour rather than the overall amount or timing of activity.

There is a growing body of literature indicating that responses to ALAN are highly variable across species and ecological contexts. While many studies have demonstrated strong behavioural or physiological responses to artificial light (e.g., [48–50], several recent studies have also reported limited responses to artificial light in some systems [51–53]. Together, these findings suggest that the ecological consequences of ALAN are unlikely to be universal and may depend strongly on species-specific traits, habitat use, behavioural ecology, and the behavioural scale at which responses are measured.

We did not detect significant effect of ALAN on predation risk. If crickets maintain similar daily activity patterns under both ALAN and control conditions, their temporal exposure to predators would likewise remain similar. Although movements in and out of the burrow were reduced under ALAN, individuals also spent longer outside during the day. Such a redistribution of activity across time periods may have helped balance overall predation exposure. Moreover, the effects of artificial illumination on predation risk often depend not only on prey behaviour but also on how predators respond to changes in light conditions, which can alter predator–prey interactions rather than affecting only one trophic level [19, 54]. For instance, insect aggregations around artificial lights can create foraging opportunities for predators such as bats and spiders [55–57]. Increased visibility caused by artificial lighting is often assumed to favour predators by making prey more conspicuous [54], and elevated predation risk under brighter conditions has been documented in several systems [58–60]. However, evidence for such effects in arthropod systems is mixed. For instance, predation on waxworm (*Galleria mellonella*) larvae was greater along habitat edges than in the middle of habitats during the day, but nocturnal predation rates along edges were not affected by the presence of street lighting [61]. In the case of *G. campestris*, predators such as robins, shrews, and lizards may respond weakly or inconsistently to the artificial illumination applied at cricket burrows, thus minimising potential effects on predation outcomes.

We found some evidence suggestive of an effect of ALAN on burrow abandonment, although this pattern was only weakly supported e (P = 0.09, Table 6). Individuals exposed to ALAN showed a higher tendency to abandon their burrows than controls. While this result should be interpreted cautiously, it raises the possibility that artificial illumination may alter refuge-use decisions or increase the costs associated with maintaining or relocating burrows. For a burrow-dependent species, such behavioural changes could carry downstream consequences, as additional investment in burrow searching or excavation may trade off against other functions such as self-maintenance, predator avoidance, growth, or future reproduction.

We also observed clear sex differences in activity timing and duration. Males exhibited earlier activity onset and later activity cessation compared to females, resulting in a longer daily activity windows. In addition, males spent significantly more time being active during the daytime window. These patterns are consistent with the behavioural ecology of field crickets. In adult *G. campestris*, males invest substantial time in mate attraction and competition, including calling and territorial interactions around their burrows [62–64]. Although the individuals examined here were nymphs, early behavioural differences between sexes may be associated with behavioural strategies that later become more pronounced during adulthood [65]. Increased activity outside the burrow may therefore reflect the trade-off between increased opportunities for resource acquisition or future reproductive opportunities and increased exposure to predation risk [66].

An additional consideration is that responses to environmental change may be more detectable in fine-scale behavioural metrics than in broader measures commonly used in previous studies. Subtle effects under natural conditions can be missed when analyses focus only on, for example, activity onset, cessation, or total activity duration.

By combining continuous monitoring with AI-assisted behavioural extraction, we quantified large numbers of individual ‘out’ events and their durations at a temporal resolution that would be highly time-consuming to obtain manually. This approach revealed treatment differences in outing behaviour that were not apparent from broader activity measures. Our results therefore highlight the value of high-resolution automated tracking for detecting subtle behavioural responses in animals in natural systems.

In summary, our results suggest that relatively dim artificial illumination similar to that produced by street lighting may have limited effects on activity timing, activity duration, and several fitness-related behaviours in *Gryllus campestris* under natural field conditions. However, ALAN can still modify fine-scale behaviour, particularly by reducing the frequency of exit from burrow. Artificial light at night is a rapidly expanding environmental pressure, and understanding how species respond under natural ecological conditions remains a major challenge [2]. Future experimental studies that combine long-term field monitoring with more detailed behavioural tracking, controlled illumination experiments and predator activity measurements will be particularly valuable for identifying when and how ALAN influences insect behaviour and fitness in natural populations.

## Acknowledgments

All authors have no conflict of interests to declare. We sincerely thank Richard Everson for contributing Python code that was instrumental in developing the GryllAI pipeline, Erik Postma for his valuable advice on the analyses, and Mark Pitt for his assistance with the experimental setup in the field. The GryllAI system was conceived and developed by R.L. and T.O.-W.; both R.L. and T.O.-W. contributed to its design, testing, and refinement. This research was supported by a Natural Environment Research Council grant (NE/V000772/1). The LED lighting equipment used in this study was supported by the University of Glasgow. R.L. was supported by the China Scholarship Council.

## Data accessibility statement

The dataset used for statistical analyses will be deposited in a public repository upon acceptance of the manuscript. The GryllAI scripts used in this study are available from GitHub: https://github.com/Ruonan325/GryllAI.

