## Supplementary Material for "Deep learning reveals the individual-level effects of artificial light at night on a wild insect"

**Table S1.** Summary of evaluation metrics for individual models.

| Model | Precision | Recall | F1 Score | Top-1 Accuracy |
|---|---|---|---|---|
| **Cricket Detection** | 0.971 | 0.985 | 0.978 | - |
| **Entrance ID** | 0.993 | 0.990 | 0.992 | - |
| **Orientation** | 0.918 | 0.949 | 0.934 | - |
| **Burrow Status** | - | - | - | 0.975 |
| **Light Level** | - | - | - | 1.000 |

Commented [TT1]: Needs explanation for what these are - this could go in the table legend

Commented [TO2R1]: I have now added explanations to the text where these metrics are first mentioned.

**Table S2.** Summary of intraclass correlation coefficient analyses comparing automated and manually-derived results.

| Factor | Intraclass Correlation Coefficient | df | *P* |
|---|---|---|---|
| **Activity Duration** | 0.917 | 59 | <0.001 |
| **BAT and EAT** | 0.999 | 33 | <0.001 |
| **Event Count** | 0.675 | 39 | <0.001 |
| **Event Duration** | 0.898 | 39 | <0.001 |

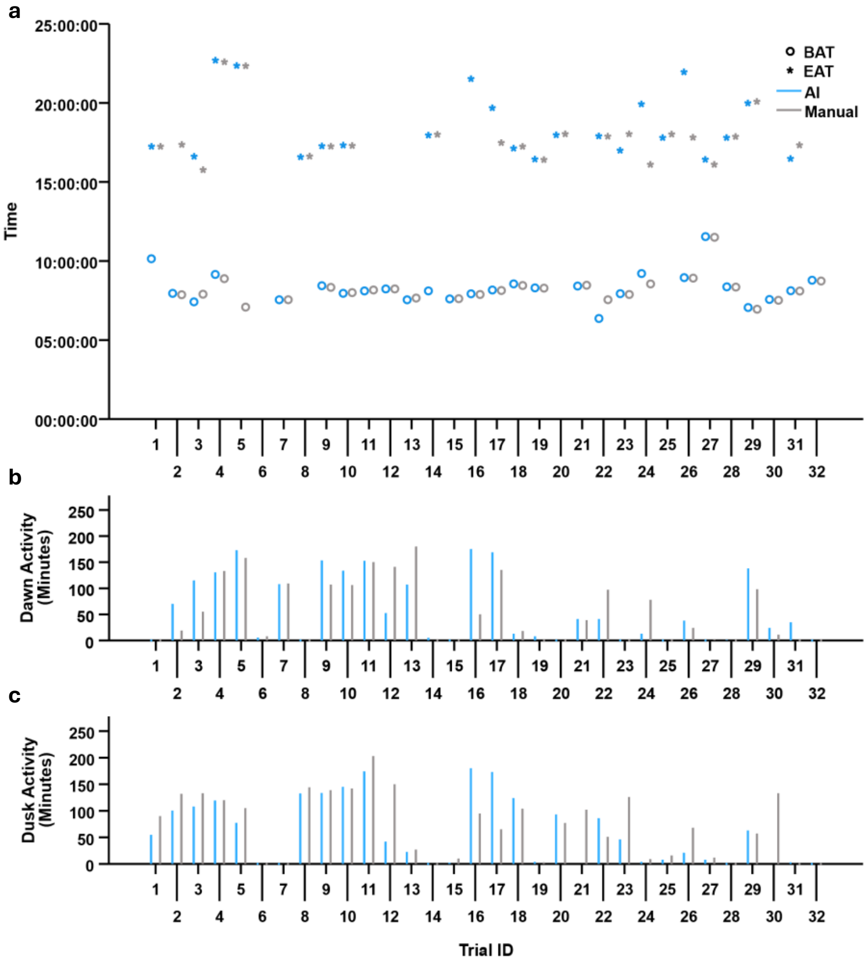


**Figure S1.** Automated and manually-derived results across system validation trials for onset of activity (BAT) and cessation of activity (EAT) timings, along with total activity durations (*N*=32). Each trial encompasses summary data for a single day and burrow, detailing (a) BAT and EAT values; (b) total activity durations during dawn (03:00–06:00); and (c) total activity durations during dusk (19:00–22:00). Colours indicate whether the data was collected manually by a human observer, or automatically using the GryllAI system (AI, blue; manual, grey).

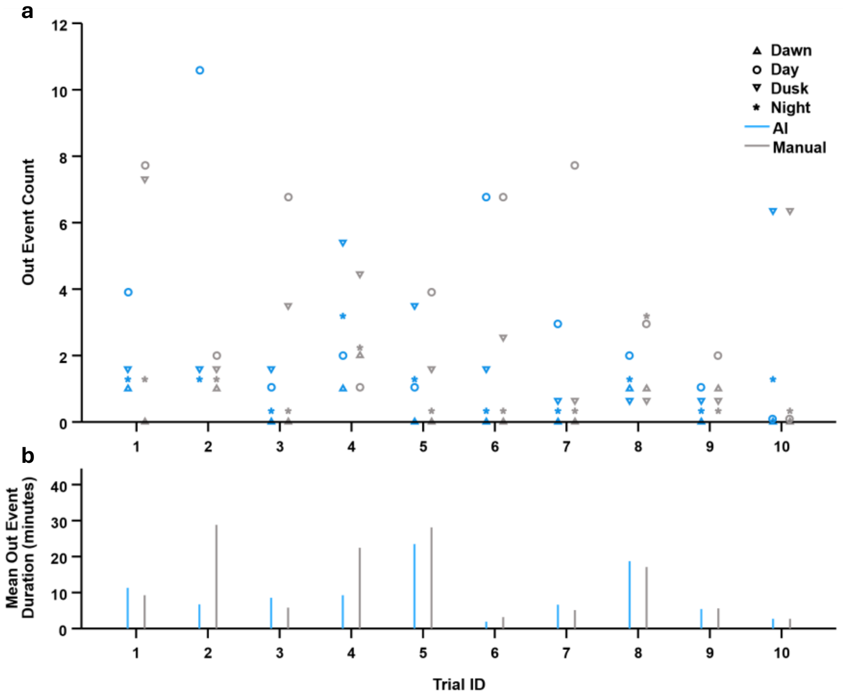


**Figure S2.** Automated and manually-derived results across system validation trials for out event counts and durations (*N*=10). Each trial encompasses summary data for a single day and burrow, detailing (a) out event counts across each time window; and (b) mean out event durations. Point shapes indicate time windows (dawn, triangle; day, circle; dusk, inverted triangle; night, asterisk), and colours indicate whether the data was collected manually by a human observer, or automatically using the GryllAI system (AI, blue; manual, grey).

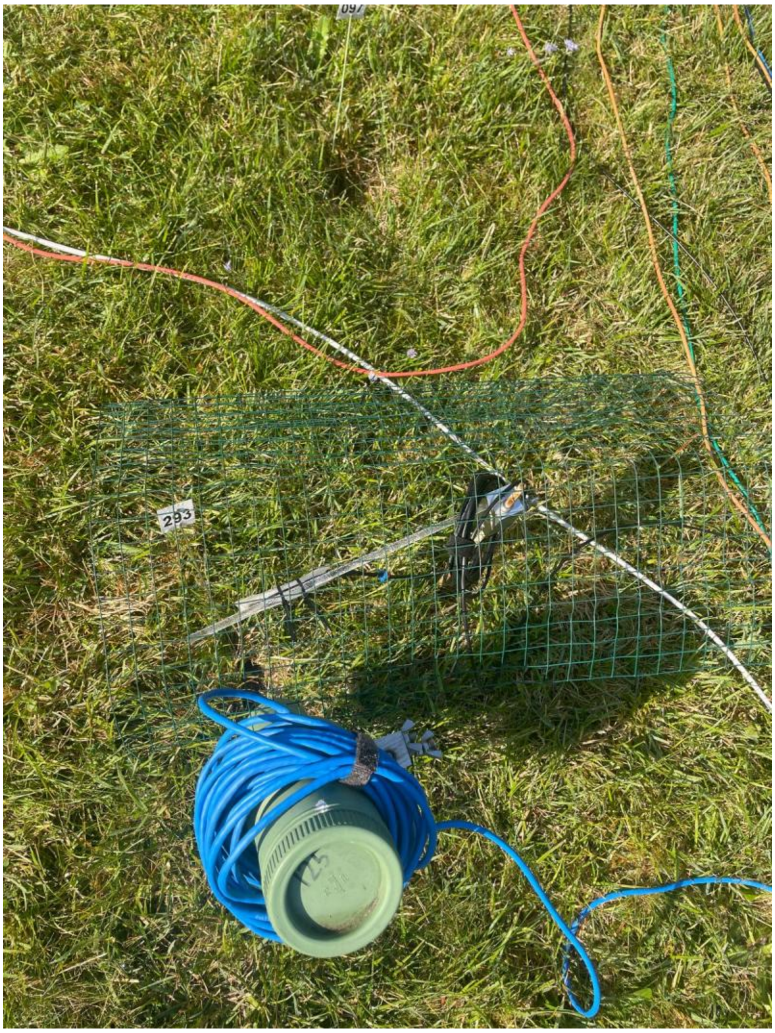

**Figure S3.** Experimental light, camera, and mesh setup positioned above a burrow.

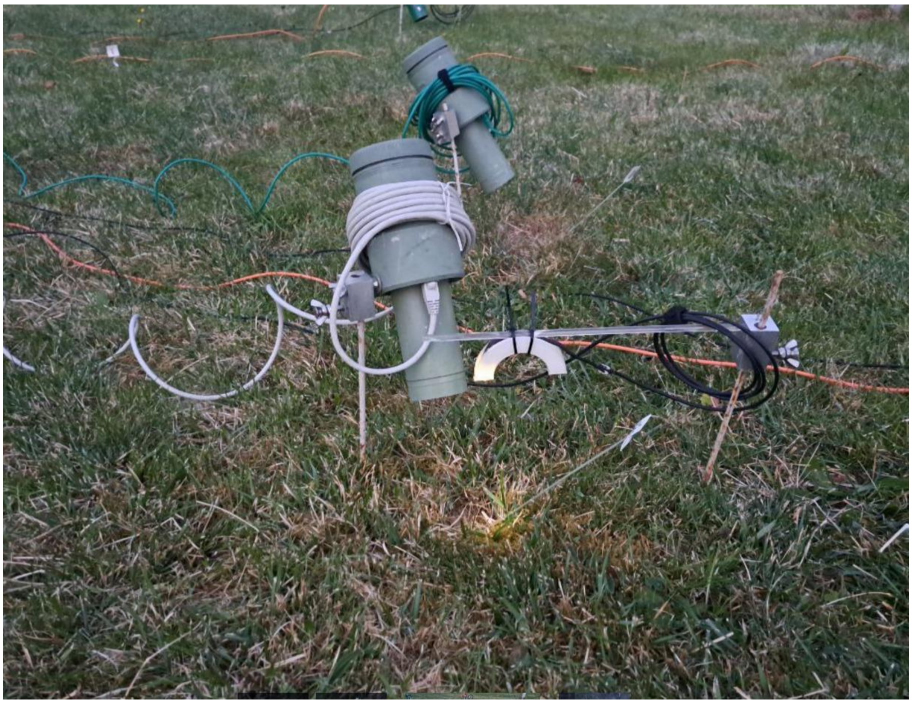

**Figure S4.** Experimental light conditions at dusk, shown without the mesh covering the burrow for clarity.

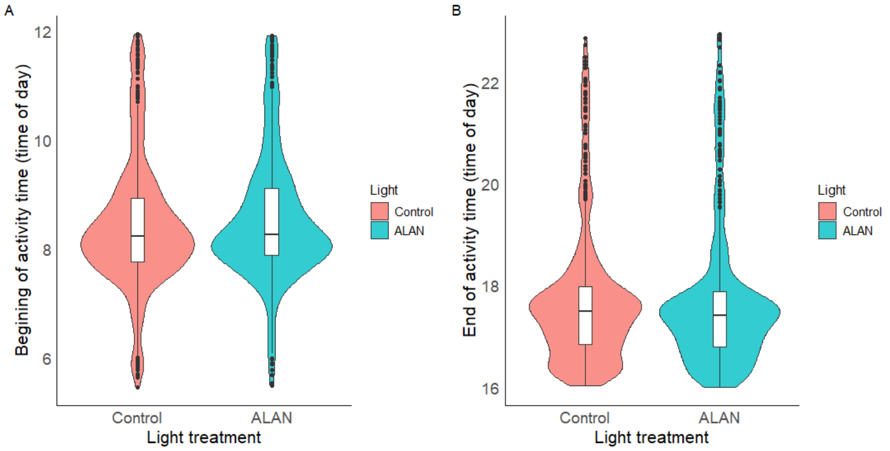

**Figure S5.** Distribution of beginning (A) and end (B) of activity time under control and ALAN light treatments. Violin plots represent the distribution density of the data, with embedded boxplots indicating the median and interquartile range. Points represent outliers beyond 1.5 × interquartile range.

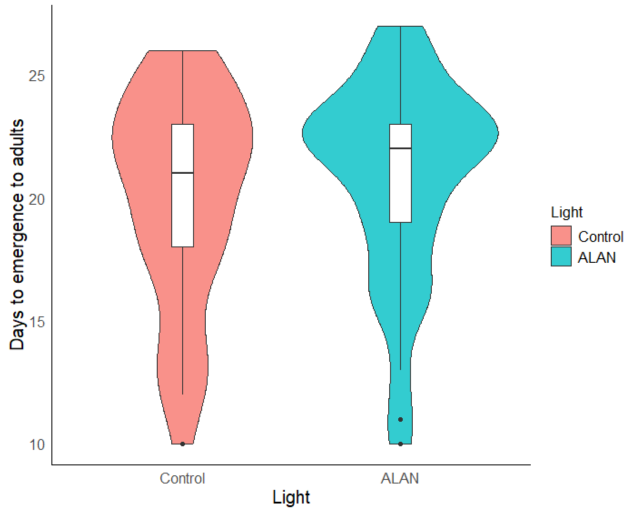


**Figure S6.** Days to emergence to adulthood under control and the artificial light at night treatments. Distribution of the days to emergence under control and ALAN conditions. Violin plots represent the distribution density of the data, with embedded boxplots indicating the median and interquartile range. Points represent outliers beyond 1.5 × interquartile range.
